# Learning place cells, grid cells and invariances: A unifying model

**DOI:** 10.1101/102525

**Authors:** Simon N. Weber, Henning Sprekeler

## Abstract

Neurons in the hippocampus and adjacent brain areas show a large diversity in their tuning to location and head direction. The underlying circuit mechanisms are not fully resolved. In particular, it is unclear why certain cell types are selective to one spatial variable, but invariant to another. For example, a place cell is highly selective to location, but typically invariant to head direction. Here, we propose that all observed spatial tuning patterns – in both their selectivity and their invariance – are a consequence of the same mechanism: Excitatory and inhibitory synaptic plasticity that is driven by the spatial tuning statistics of synaptic inputs. Using simulations and a mathematical analysis, we show that combined excitatory and inhibitory plasticity can lead to localized, grid-like or invariant activity. Combinations of different input statistics along different spatial dimensions reproduce all major spatial tuning patterns observed in rodents. The model is robust to changes in parameters, develops patterns on behavioral time scales and makes distinctive experimental predictions. Our results suggest that the interaction of excitatory and inhibitory plasticity is a general principle for the formation of neural representations.

---

Neurons in the hippocampus and the adjacent regions exhibit a broad variety of spatial activation patterns that are tuned to position, head direction or both. Common observations in these spatial dimensions are localized, bell shaped tuning curves [1, 2], periodically repeating activity [3, 4] and invariances [5, 6], as well as combinations of these along different spatial dimensions [7, 8]. For example, head direction cells are often invariant to location [6], and place cells are commonly invariant to head direction [5].

The cellular and network mechanisms that give rise to each of these firing patterns are subject to extensive experimental and theoretical research. Several computational models have been suggested to explain the emergence of grid cells [9–21], place cells [11, 22–27] and head direction cells [11, 28–30].

Most of these models are designed to explain the spatial selectivity of one particular cell type and do not consider invariances along other dimensions, although the formation of invariant representations is a non-trivial problem [31].

In view of the variety of spatial tuning patterns, the question arises if the differences in tuning of different cells in different areas reflect differences in microcircuit connectivity, single cell properties or plasticity rules, or if there is a unifying principle. In this paper we suggest that both the observed spatial selectivities and invariances can be explained by a common mechanism – interacting excitatory and inhibitory synaptic plasticity – and that the observed differences in the response profiles of grid, place and head direction cells result from differences in the spatial tuning of excitatory and inhibitory synaptic afferents. Here, we explore this hypothesis in a computational model of a feedforward network of rate-based neurons. Simulations as well as a mathematical analysis indicate that the model reproduces the large variety of response patterns of neurons in the hippocampal formation and adjacent areas and make predictions for the input statistics of each cell type.

The suggested mechanism ports the robust pattern formation of attractor models [9, 10, 12] from the neural to the spatial domain and increases the speed of selforganization of plasticity-based mechanisms [15, 17–19] to time scales on which the spatial tuning of neurons is typically measured.

## Results

We study the development of spatial representations in a network of rate-based neurons with interacting excitatory and inhibitory plasticity. A single model neuron that represents a cell in the hippocampal formation or adjacent areas receives feedforward input from excitatory and inhibitory synaptic afferents. As a simulated rat moves through an environment, these synaptic afferents are weakly modulated by spatial location and in later sections also by head direction. This modulation is irregular and non-localized with multiple maxima (Fig. 1a, Methods) [32]. Importantly, different inputs show different modulation profiles. We also show results for localized, i.e., place cell-like, input [33–35]. The output rate is given by a weighted sum of the excitatory and inhibitory inputs.

In our model, both excitatory and inhibitory synaptic weights are subject to plasticity. The excitatory weights change according to a Hebbian plasticity rule [36] that potentiates the weights in response to simultaneous pre-and postsynaptic activity. The inhibitory synapses evolve according to a plasticity rule that changes their weights in proportion to presynaptic activity and the difference between postsynaptic activity and a target rate (1 Hz in all simulations). This rule has previously been shown to balance excitation and inhibition such that the firing rate of the output neuron approaches the target rate [37]. We assume the inhibitory plasticity to act fast enough to track changes of excitatory weights, so that excitation and inhibition are approximately balanced at all times.

### The relative spatial smoothness of the excitatory and inhibitory input determines the firing pattern of the output neuron

We first simulate a rat that explores a linear track (Fig. 1). The spatial tuning of each input neuron is random but depends smoothly on the location of the animal (e.g., Fig. 1a). As a measure of smoothness, we use the spatial autocorrelation length of the inputs. In the following, this is the central parameter of the input statistics, which is chosen separately for excitation and inhibition.

**Figure 1:**
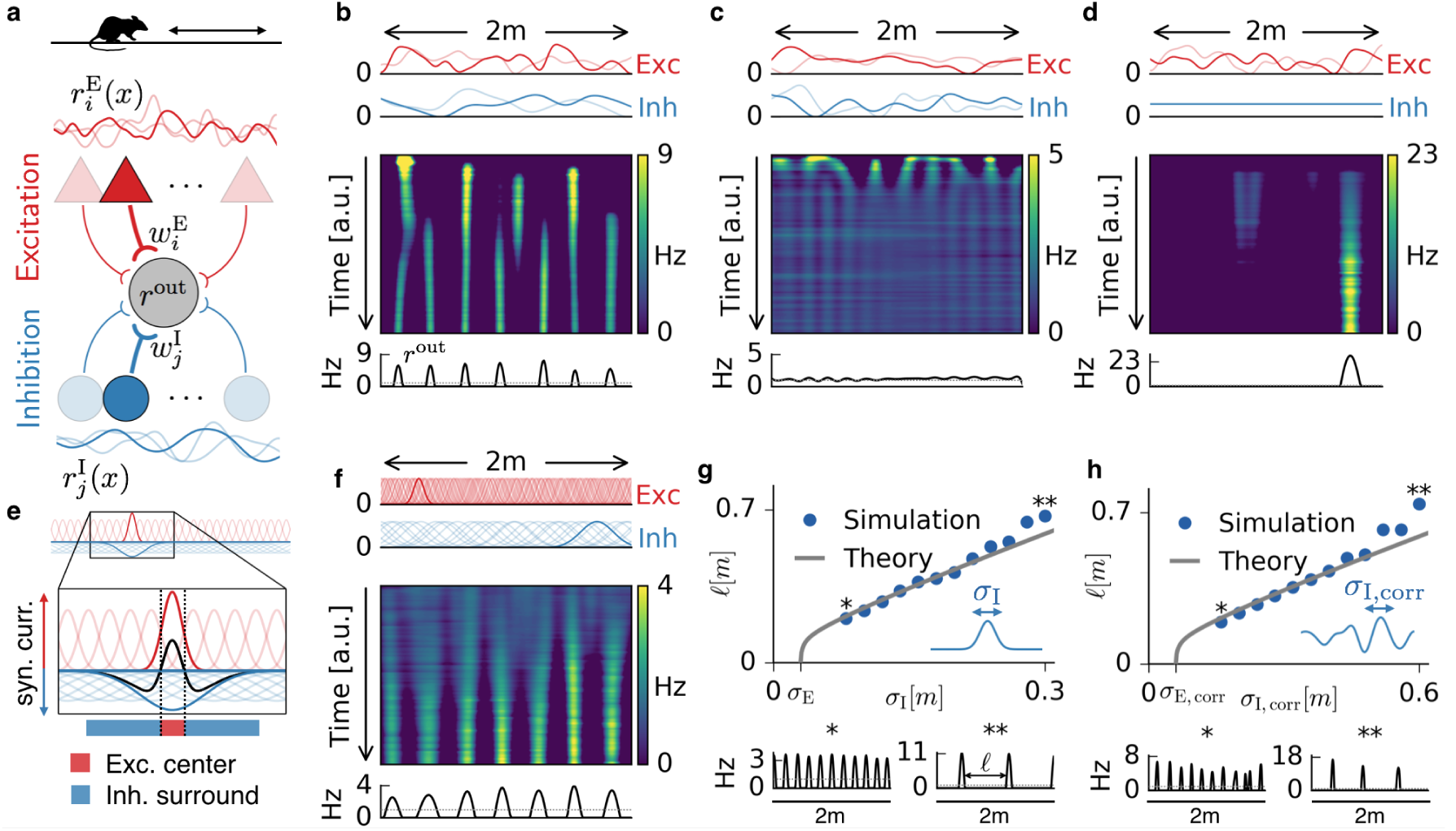
Emergence of periodic, invariant and single field firing patterns. **a**) Network model for a linear track. A threshold-linear output neuron (gray) receives input from excitatory (red) and inhibitory (blue) cells, which are spatially tuned (curves on top and bottom). **b**) Spatially tuned input with smoother inhibition than excitation. The fluctuating curves (top) show two exemplary spatial tunings (one is highlighted) of excitatory and inhibitory input neurons. Interacting excitatory and inhibitory synaptic plasticity gradually changes an initially random response of the output neuron (firing rate *r*^out^) into a periodic, grid cell-like activity pattern. **c**) If the spatial tuning of inhibitory input neurons is less smooth than that of excitatory input neurons the interacting excitatory and inhibitory plasticity leads to a spatially invariant firing pattern. The output neuron fires close to the target rate of 1 Hz everywhere. **d**) For very smooth or spatially untuned inhibitory inputs, the output neuron develops a single firing field, reminiscent of a place cell. **e**) The mechanism, illustrated for place cell-like input. When a single excitatory weight is increased relative to the others, the balancing inhibitory plasticity rule leads to an immediate increase of inhibition at the associated location. If inhibitory inputs are smoother than excitatory inputs, the resulting approximate balance creates a center surround field: a local overshoot of excitation (firing field) surrounded by an inhibitory corona. The next firing field emerges at a distance where the inhibition has faded out. Iterated, this results in a spatially periodic arrangement of firing fields. **f**) Inputs with place field-like tuning. Gaussian curves (top) show the spatial tuning of excitatory and inhibitory input neurons (one neuron of each kind is highlighted, 20 percent of all inputs are displayed). A grid cell firing pattern emerges from an initially random weight configuration. **g**) Grid spacing ℓ scales with inhibitory tuning width *σ*_I_. Simulation results (dots) agree with a mathematical bifurcation analysis (solid). Output firing rate examples at the two indicated locations are shown at the bottom. **h**) Inhibitory smoothness *σ*_I_,_corr_ controls grid spacing; arrangement as in **d**.

At the beginning of each simulation, all synaptic weights are random. As the animal explores the track, the excitatory and inhibitory weights change in response to pre- and postsynaptic activity, and the output cell gradually develops a spatial activity pattern. We find that this pattern is primarily determined by whether the excitatory or inhibitory inputs are smoother in space. If the inhibitory tuning is smoother than the excitatory tuning (Fig. **1b**), the output neuron develops equidistant firing fields, reminiscent of grid cells on a linear track [38]. If instead the excitatory tuning is smoother, the output neuron fires close to the target rate of 1 Hz everywhere (Fig. **1c**); it develops a spatial invariance. For spatially untuned inhibitory afferents, the output neuron develops a single firing field, reminiscent of a one-dimensional place cell (Fig. 1d, compare [39]).

The emergence of these firing patterns can be best explained in the simplified scenario of place field-like input tuning **(Fig. 1e,f**). The spatial smoothness is then given by the size of the place fields. Let us assume that the output neuron fires at the target rate everywhere (Supplementary Online Material, SOM). From this homogeneous state, a small potentiation of one excitatory weight leads to an increased firing rate of the output neuron at the location of the associated place field (highlighted red curve in Fig. 1e). To bring the output neuron back to the target rate, the inhibitory learning rule increases the synaptic weight of inhibitory inputs that are tuned to the same location (highlighted blue curve in Fig. 1e). If these inhibitory inputs have smaller place fields than the excitatory inputs (Fig. 1c), this restores the target rate everywhere [37]. Hence, inhibitory plasticity can stabilize spatial invariance if the inhibitory inputs are sufficiently precise (i.e., not too smooth) in space. In contrast, if the spatial tuning of the inhibitory inputs is smoother than that of the excitatory inputs, the target firing rate cannot be restored everywhere. Instead, a compensatory potentiation of inhibitory weights increases the inhibition in a spatial region that has at least the size of the inhibitory place fields. This leads to a corona of inhibition, in which the output neuron cannot fire (Fig. 1e, blue region). Outside of this inhibitory surround the output neuron can fire again and the next firing field develops. Iterated, this results in a periodic arrangement of firing fields (Fig. 1f). Spatially untuned inhibition corresponds to an infinitely large inhibitory corona that exceeds the length of the linear track, so that only a single place field remains.

The argument of the preceding paragraph can be extended to the scenario where input is only weakly modulated by space. For non-localized input tuning **(Fig. 1b,c,d**), any weight change that increases synaptic input in one location will also increase it in a surround that is given by the smoothness of the input tuning (see SOM for a mathematical analysis). In the simulations, the randomness manifests itself in occasional defects in the emerging firing pattern (Fig. 1h, bottom). The above reasoning suggests that the width of individual firing fields is determined by the smoothness of the excitatory input tuning, while the distance between grid fields, i.e., the grid spacing, is set by the smoothness of the inhibitory input tuning. Indeed, both simulations and a mathematical analysis (SOM) confirm that the grid spacing scales linearly with the inhibitory smoothness in a large range, both for localized (Fig. 1g) and non-localized input tuning (Fig. 1h). In summary, the interaction of excitatory and inhibitory plasticity can lead to spatial invariance, spatially periodic activity patterns or single place fields depending on the spatial statistics of the excitatory and inhibitory input.

### Emergence of hexagonal firing patterns

When a rat navigates in a two-dimensional arena, the spatial firing maps of grid cells in the medial entorhinal cortex (mEC) show a pronounced hexagonal symmetry [3, 4] with different grid spacings and spatial phases. To study whether a hexagonal firing pattern can emerge from interacting excitatory and inhibitory plasticity, we simulate a rat in a quadratic box. The rat explores the arena for 10 hours, following trajectories extracted from behavioral data [40] (SOM). To investigate the role of the input statistics, we consider three different classes of input tuning: i) place cell-like input (Fig. 2a), ii) sparse non-localized input, in which the tuning of each input neuron is given by the sum of 100 randomly located place fields (Fig. 2b) and iii) dense non-localized input, in which the tuning of each input is a random function with fixed spatial smoothness (Fig. 2c). For all input classes, the spatial tuning of the inhibitory inputs is smoother than that of the excitatory inputs.

**Figure 2:**
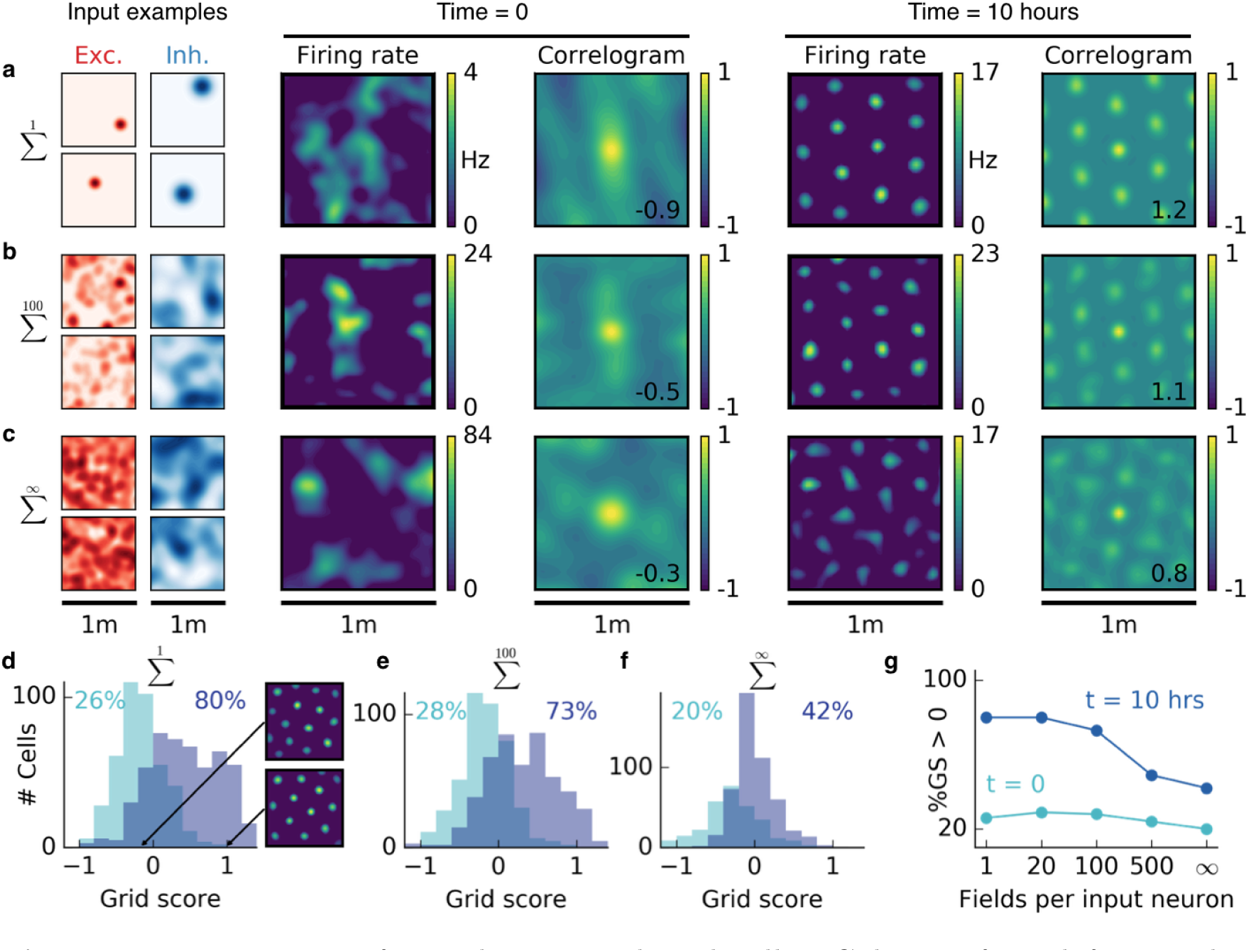
Emergence of two dimensional grid cells. Columns from left to right: Spatial tuning of excitatory and inhibitory input neurons (two examples each); spatial firing rate map of the output neuron and corresponding autocorrelogram before and after spatial exploration of 10 hours. The number on the correlogram shows the associated grid score. Different rows correspond to different spatial tuning characteristics of the excitatory and inhibitory input. For all figures the spatial tuning of inhibitory input neurons is smoother than that of excitatory input neurons. **a**) Each input neuron is a place cell with random location. **b**) The tuning of each input neuron is given as the sum of 100 randomly located place fields. **c**) The tuning of each input neuron is a random smooth function of the location. This corresponds to the sum of infinitely many randomly located place fields. Before learning the spatial tuning of the output neuron shows no symmetry. After 10 hours of spatial exploration the output neuron developed a hexagonal pattern. **d**) Grid score histogram for 500 output cells with place cell-like input. Before learning (light blue) 26% of the output cells have a positive grid score. After 10 hours of spatial exploration this value increases to 80%. Two example rate maps are shown. The arrows point to the grid score of the associated rate map. Even for low grid scores the learned firing pattern looks grid-like. **e,f**) Grid score histograms for input tuning as in **b,c**, arranged as in **d**. **g**) Fraction of neurons with positive grid score before (light blue) and after learning (dark blue) as a function of the number of fields per input neuron.

Initially, all synaptic weights are random and the activity of the output neuron shows no spatial symmetry. While the rat forages through the environment, the output cell develops a periodic firing pattern for all three input classes, reminiscent of grid cells in the mEC [3, 4] and typically with the same hexagonal symmetry. This hexagonal arrangement is again due to the smoother inhibitory input tuning, which generates a spherical inhibitory corona around each firing field (compare Fig. 1e). These center-surround fields arrange in a hexagonal pattern – the closest packing of spheres in two dimensions; compare [41]. We find that the spacing of this pattern is determined by the inhibitory smoothness, whereas the orientation and the phase of the grid depend in decreasing order on the random initialization of the input tuning, the trajectories and the initial synaptic weights (SOM, Fig. **S9**).

For the linear track, the randomness of the non-localized inputs leads to defects in the periodicity of the grid pattern. In two dimensions, we find that the randomness leads to distortions of the hexagonal grid. To quantify this effect, we simulated 500 random trials for each of the three input scenarios and plotted the grid score histogram (SOM) before and after 10 hours of spatial exploration **(Fig. 2d,e,f**). Different trials have different trajectories, different initial synaptic weights and different random locations of the input place fields (for sparse input) or different random input functions (for dense input). For place cell-like input, most of the output cells develop a positive grid score during 10 hours of spatial exploration (26% before to 80% after learning, Fig. 2d). Even for low grid scores, the firing rate maps look grid-like after learning but exhibit a distorted symmetry (Fig. 2d). For sparse non-localized input, the fraction of output cells with a positive grid score increases from 28% to 73% and for dense non-localized input from 20% to 42% within 10 hours of spatial exploration.

In summary, the interaction of excitatory and inhibitory plasticity leads to gridlike firing patterns in the output neuron for all three input scenarios. The grids are typically less distorted for sparser input (Fig. 2g).

### Rapid appearance of grid cells

In unfamiliar environments, neurons in the mEC exhibit grid-like firing patterns within minutes [4]. Moreover, grid cells react quickly to changes in the environment [42–44]. These observations challenge models for grid cells that require gradual synaptic changes during spatial exploration. In principle, the time scale of plasticity-based models can be augmented arbitrarily by increasing the synaptic learning rates. For stable patterns to emerge, however, significant weight changes must occur only after the animal has visited most of the environment. To explore the edge of this trade-off between speed and stability, we increased the learning rates to a point where the grids are still stable but where further increase would reduce the stability (SOM, Fig. **S8**). For place cell-like input, periodic patterns can be discerned within 10 minutes of spatial exploration, starting with random initial weights **(Fig. 3a,b**). The pattern further emphasizes over time and remains stable for many hours (Fig. **3c**).

**Figure 3:**
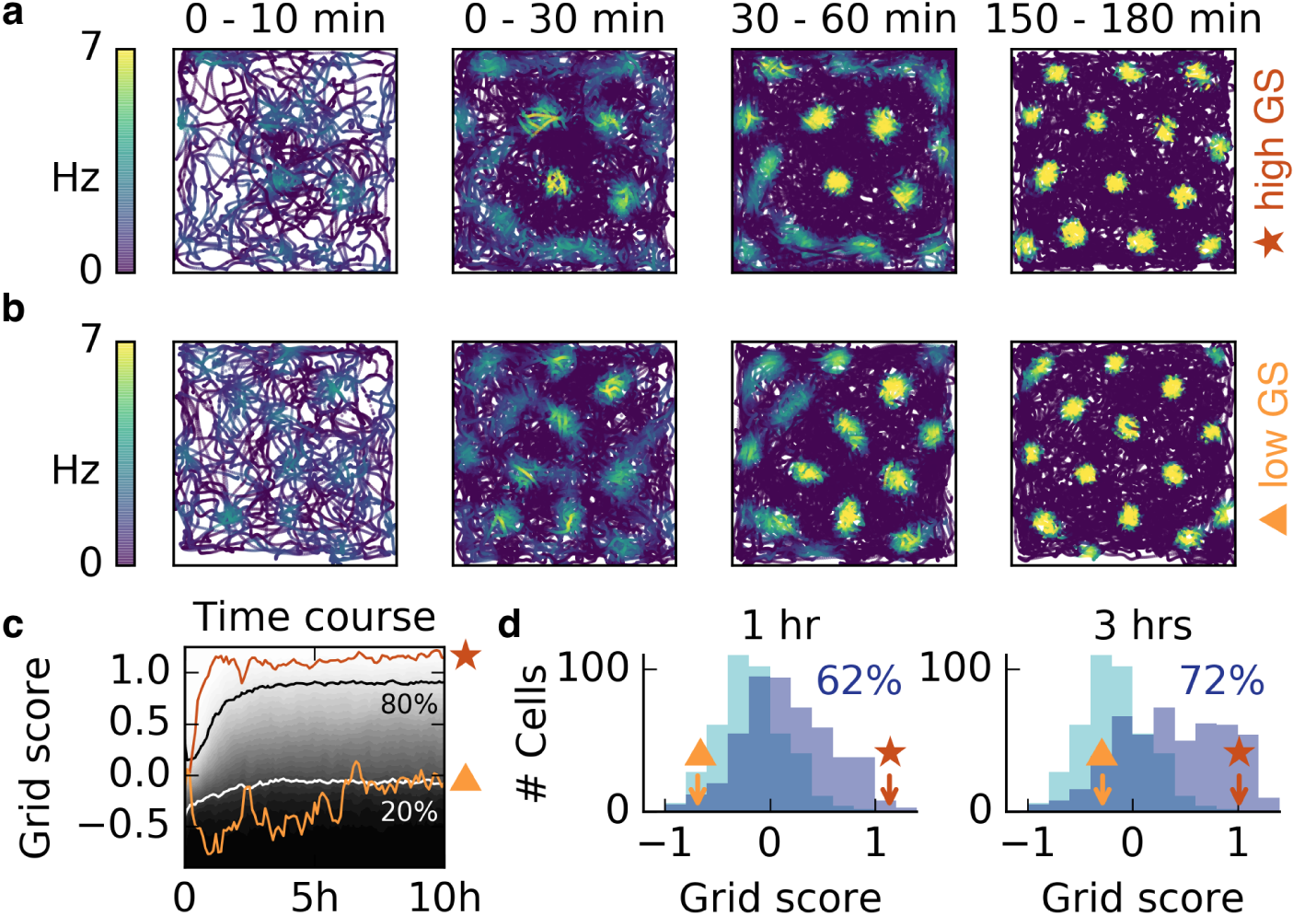
Grid patterns form rapidly during exploration and remain stable for many hours. **a,b**) Rat trajectories with color-coded firing rate. Bright colors indicate higher firing rates. Initially all synaptic weights are set to random values. Part **a** and **b** show two different realizations with a good (red star) and a bad (orange triangle) grid score development. After few minutes a periodic structure becomes visible and enhances over time. **c**) Time course of the grid score in the simulations shown in **a** (red) and **b** (orange). While the periodic patterns emerge within minutes, the manifestation of the final hexagonal pattern typically takes a couple of hours. Once the pattern is established it remains stable for many hours. The gray scale shows the cumulative histogram of the grid scores of 500 realizations (black=0, white=1). The solid white and black lines indicate the 20% and 80% percentile, respectively. **d**) Histogram of grid scores of the 500 simulations shown in **c**. Initial histogram in light blue, histogram after 1 hour and after 3 hours in dark blue. Numbers show the fraction of cells with positive grid score at the given time. Rat trajectories taken from [40].

To investigate the robustness of this phenomenon we ran 500 realizations with different trajectories, initial synaptic weights and locations of input place fields. In all simulations, a periodic pattern emerged within the first 30 minutes, and a majority of patterns exhibited hexagonal symmetry after three hours (increasing from 26% to 72%,**Fig. 3c,d**). For non-localized input, the emergence of the final grids typically takes longer, but the first grid fields are also observed within minutes and are still present in the final grid, as observed in experiments [4] (SOM, Fig. **S6**).

In summary, periodic patterns emerge rapidly in our model and the associated time scale is limited primarily by how quickly the animal visits its surroundings, i.e., by the same time scale that limits the experimental recognition of the grids.

### Place cells, band cells and stretched grids

In addition to grids, the mEC and adjacent brain areas exhibit a plethora of other spatial activity patterns including spatially invariant [6], band-like [8] (periodic along one direction and invariant along the other), and spatially periodic but non-hexagonal patterns [8]. Moreover, place cells in the hippocampus proper are typically only tuned to a single or few locations in a given environment [33, 45, 46]. If the animal traversed the environment along a straight line, all of these cells would be classified as periodic, localized or invariant (Fig. 1), although the classification could vary depending on the direction of the line. Based on this observation, we hypothesized that all of these patterns could be the result of an input autocorrelation structure that differs along different spatial directions.

We first verified that also in a two-dimensional arena, place cells emerge from a very smooth inhibitory input tuning (Fig. 4a) and spatial invariance results when excitation is broader than inhibition (Fig. 4b). We then varied the smoothness of the inhibitory inputs independently along two spatial directions. If the spatial tuning of inhibitory inputs is smoother than the tuning of the excitatory inputs along one dimension but less smooth along the other, the output neuron develops band cell-like firing patterns (Fig. 4c). If inhibitory input is smoother than excitatory input, but not isotropic, the output cell develops stretched grids with different spacing along two axes (Fig. 4d). For these anisotropic cases, stretched hexagonal grids and rectangular arrangements of firing fields appear similarly favorable (compare Fig. 4d, second row and column). A hexagonal arrangement could be favored by a dense packing of inhibitory coronas, whereas a rectangular arrangement would maximize the proximity of the excitatory centers, given the inhibitory corona (SOM, Fig. **S10**). In summary, the relative spatial smoothness of inhibitory and excitatory input determines the symmetry of the spatial firing pattern of the output neuron. The requirements for the input tuning that support invariance, periodicity and localization apply individually to each spatial dimension, opening up a combinatorial variety of spatial tuning patterns.

**Figure 4:**
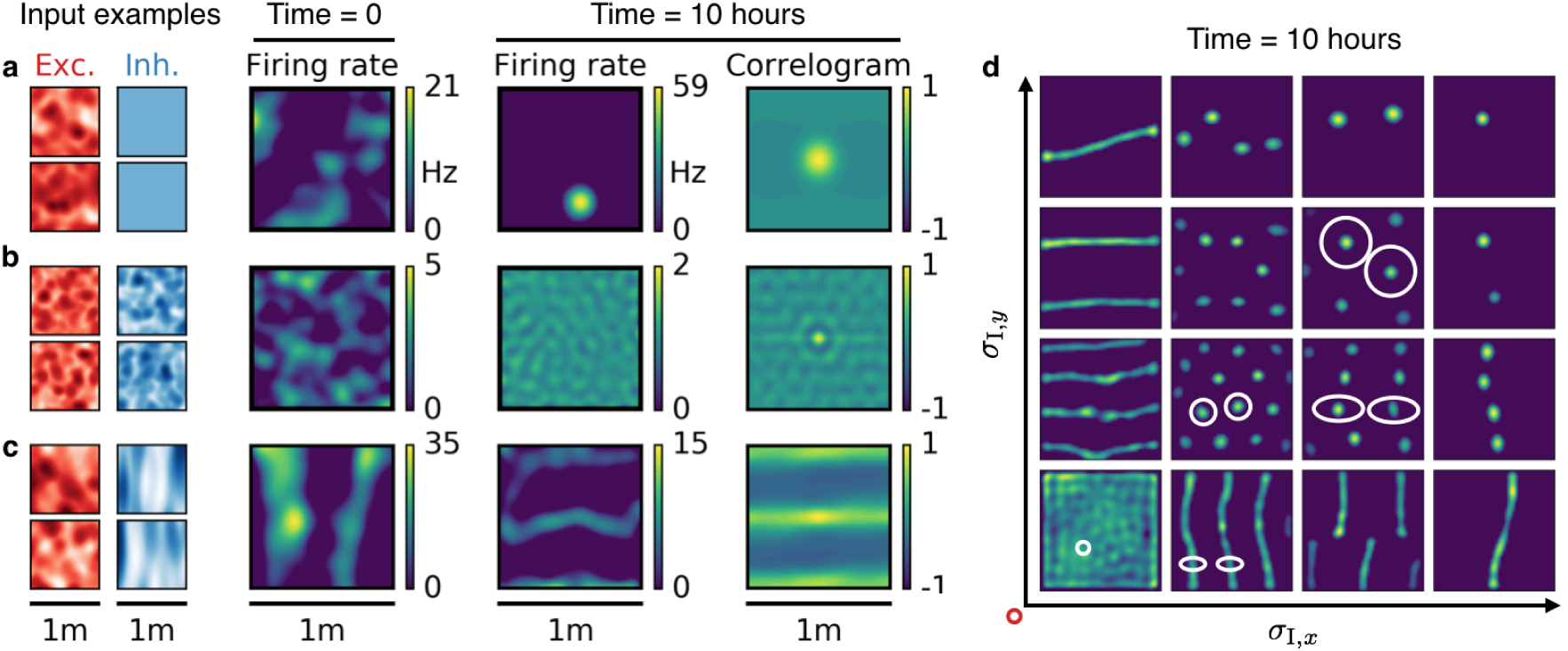
Emergence of spatially tuned cells of diverse symmetries. **a**) Place cells emerge if the inhibitory autocorrelation length exceeds the box length or if the inhibitory neurons are spatially untuned. **b**) The output neuron develops an invariance, if the spatial tuning of inhibitory input neurons is less smooth than the tuning of excitatory input neurons. **c**) Band cells emerge if the spatial tuning of inhibitory input is asymmetric, such that the autocorrelation length is larger than that of excitatory input along one direction (here the *y*-direction) and smaller along the other (here the *x*-direction). **d**) Overview of how the shape of the inhibitory input tuning determines the firing pattern of the output neuron. Each element depicts the firing rate map of the output neuron after 10 hours. White ellipses of width 2*σ*_I_*,_x_* and 2*σ*_I_*,_y_* in *x*– and *y*–direction indicate the direction-dependent standard deviation of the spatial tuning of the inhibitory input neurons. The width of the excitatory tuning fields, *σ_E_,* is the same in all simulations. The red circle at the axis origin is of diameter 2*σ_E_.*

### Spatially tuned input combined with head direction selectivity leads to grid, conjunctive and head direction cells

Many cells in and around the hippocampus are tuned to the head direction of the animal [2, 47, 48]. These head direction cells are typically tuned to a single head direction, just like place cells are typically tuned to a single location. Moreover, head direction cells are often invariant to location [6], just like place cells are commonly invariant to head direction [5]. There are also cell types with conjoined spatial and head direction tuning. Conjunctive cells in the mEC fire like grid cells in space, but only in a particular head direction [7], and many place cells in the hippocampus of crawling bats also exhibit a head direction tuning [49].

To investigate whether these tuning properties could also result in our model, we simulated a rat that moves in a square box, whose head direction is constrained by the direction of motion (SOM). Each input neuron is tuned to both space and head direction (e.g., Fig. 5a).

**Figure 5:**
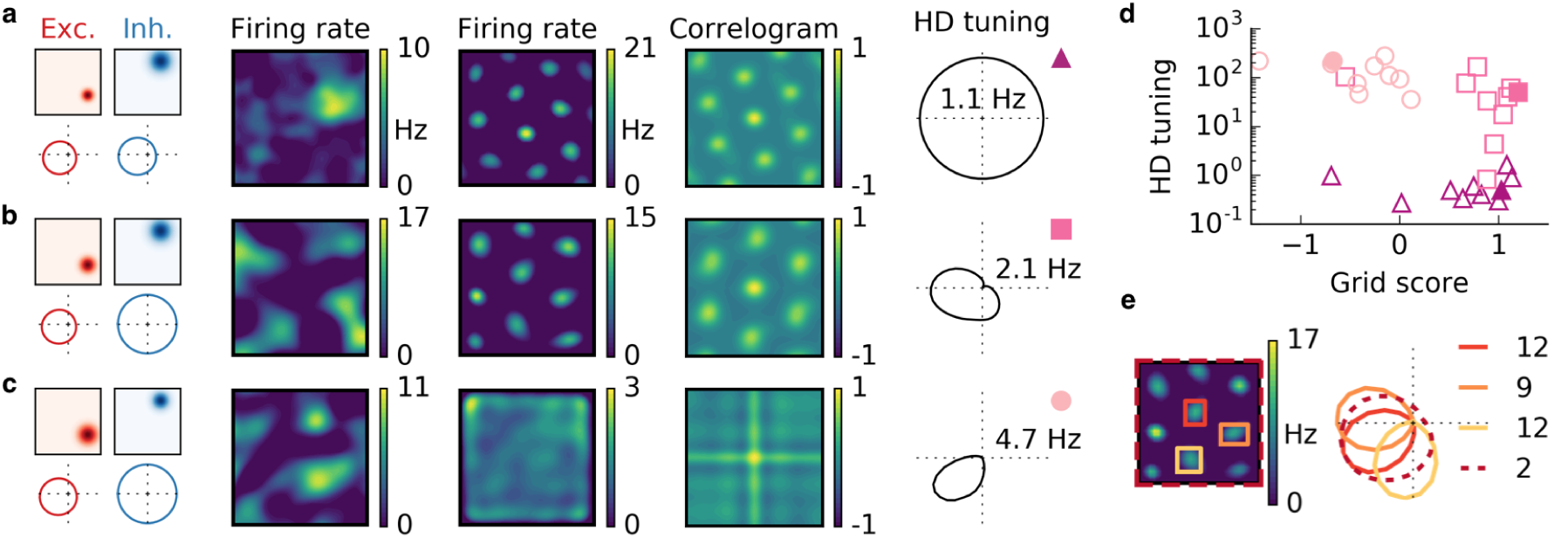
Combined spatial and head direction tuning. **a,b,c**) Columns from left to right: Spatial tuning and head direction tuning (polar plot) of excitatory and inhibitory input neurons (one example each); spatial firing rate map of the output neuron before learning and after spatial exploration of 10 hours with corresponding autocorrelogram; head direction tuning of the output neuron after learning. The numbers in the polar plots indicate the peak firing rate at the preferred head direction after averaging over space. **a**) Wider spatial tuning of inhibitory input neurons than of excitatory input neurons combined with narrower head direction tuning of inhibitory input neurons leads to a grid cell-like firing pattern in space with invariance to head direction, i.e. the output neuron fires like a pure grid cell. **b**) The same spatial input characteristics as in **a** combined with head direction-invariant inhibitory input neurons leads to grid cell-like activity in space and a preferred head direction, i.e. the output neuron fires like a conjunctive cell. **c**) If the spatial tuning of inhibitory input neurons is less smooth than that of excitatory neurons and the concurrent head direction tuning is wider for inhibitory than for excitatory neurons, the output neuron is not tuned to space but to a single head direction, i.e. the output neuron fires like a pure head direction cell. **d**) Head direction tuning and grid score of 10 simulations of the three cell types. Each symbol represents one realization with random input tuning. The markers correspond to the tuning properties of the input neurons as depicted in **a, b, c**: grid cell (triangles), conjunctive cell (squares), head direction cell (circles). The values that correspond to the output cells in **a, b, c** are shown as filled symbols. **e**) In our model, the head direction tuning of individual grid fields is sharper than the overall head direction tuning of the conjunctive cell. Depicted is a rate map of a conjunctive cell (left) and the corresponding head direction tuning (right, dashed). For three individual grid fields, indicated with colored squares, the head direction tuning is shown in the same polar plot. The overall tuning of the grid cell (dashed) is a superposition of the tuning of all grid fields. Numbers indicate the peak firing rate (in Hz) averaged individually within each of the four rectangles in the rate map. We obtained similar results for non-localized spatial and head direction input (SOM, Fig. **S7**).

In line with the previous observations, we find that the *spatial* tuning of the output neuron is determined by the relative *spatial* smoothness of the excitatory and inhibitory inputs and the *head direction* tuning of the output neuron is determined by the relative smoothness of the *head direction* tuning of the inputs from the two populations. If the head direction tuning of excitatory input neurons is smoother than that of inhibitory input neurons, the output neuron becomes invariant to head direction (Fig. 5a). If instead only the excitatory input is tuned to head direction, the output neuron develops a single activity bump at a particular head direction **(Fig. 5b,c**). The concurrent spatial tuning of the inhibitory input neurons determines the spatial tuning of the output neuron. For spatially smooth inhibitory input, the output neuron develops a hexagonal firing pattern **(Fig. 5a,b**) and for less smooth inhibitory input the firing of the output neuron is invariant to the location of the animal (Fig. 5c). In summary, the relative smoothness of inhibitory and excitatory input neurons in space and in head direction determines whether the output cell fires like a pure grid cell, a conjunctive cell or a pure head direction cell (Fig. 5d).

We find that the overall head direction tuning of conjunctive cells is broader than that of individual grid fields (Fig. 5e). This results from variations in the preferred head direction of different grid fields. Typically, however, these variations remain small enough to preserve an overall head direction tuning of the cell, because individual grid fields tend to align their head direction tuning (SOM, Fig. **S10**). Whether a narrower head direction of individual grid fields is present also in rodents is not resolved.

## Discussion

We presented a self-organization model that reproduces the experimentally observed spatial and head direction tuning patterns in the hippocampus and adjacent brain regions. Its core mechanism is an interaction of Hebbian plasticity in excitatory synapses and homeostatic Hebbian plasticity in inhibitory synapses [37, 50]. The main prediction of the model is that the spatial autocorrelation structure of excitatory and inhibitory inputs determines – and should thus be predictable from – the output pattern of the cell. Investigations of the tuning of individual cells [51] or even synapses [52] that project to spatially tuned cells would thus be a litmus test for the proposed mechanism.

The origin of synaptic input to spatially tuned cells is not fully resolved [53]. Our model is consistent with input from sensory areas, given that the mechanism is robust to the precise properties of the input and essentially only requires temporally stable tuning in a given spatial environment. The input could also stem from within the hippocampal formation, where spatial tuning has been observed in both excitatory [1] and inhibitory [34, 35] neurons. For example, the notion that mEC neurons receive input from hippocampal place cells is supported by several studies: Place cells in the hippocampus emerge earlier during development than grid cells in the mEC [54, 55], grid cells lose their tuning pattern when the hippocampus is deactivated [56] and, finally, both the firing fields of place cells and the spacing and field size of grid cells increase along the dorso-ventral axis [57-59].

Inhibition is usually thought to arise from local interneurons (but see [60]), suggesting that spatially tuned inhibitory input to mEC neurons originates from the entorhinal cortex itself. Interneurons in mEC display a spatial tuning [32] that could be inherited from hippocampal place cells or other grid cells [13]. The broader spatial tuning required for the emergence of spatial selectivity could be established, e.g., by pooling over cells with similar tuning or through a non-linear input-output transformation in the inhibitory circuitry. If inhibitory input is indeed local, the increase in grid spacing along the dorso-ventral axis [58] suggests that the tuning of inhibitory interneurons gets smoother along this axis. For smoother tuning functions, less neurons are needed to cover the whole environment, in accordance with the decrease in interneuron density along the dorso-ventral axis [61].

The observed spatial tuning patterns have also been explained by other models. In continuous attractor networks (CAN), each cell type could emerge from a specific connectivity pattern, combined with a mechanism that translates the motion of the animal into shifts of neural activity on an attractor. How the required connectivity patterns could emerge is subject to debate [62]. A measurable distinction of our model from CAN models is its response to a global reduction of inhibition. While a modification of inhibition typically changes the grid spacing in CAN models of grid cells [13, 63], the grid field locations generally remain untouched in our model. The grid fields merely change in size, until inhibition is recovered by inhibitory plasticity (Fig. 6a). This can be understood by the colocalization of the grid fields and peaks in the excitatory membrane current **(Fig. 6b,c**). A reduction of inhibition leads to an increased protrusion of these excitatory peaks and thus to wider firing fields. The observed temporal stability of mEC grid patterns in spite of dopaminergic modulations of GABAergic transmission would be in line with our model [64]. Moreover, we found that for localized input tuning, the inhibitory membrane current typically also peaks at the locations of the grid fields. This co-tuning breaks down for non-localized input (Fig. 6b). In contrast, CAN models predict that the inhibitory membrane current has the same periodicity as the grid, but possibly phase shifted [65].

**Figure 6:**
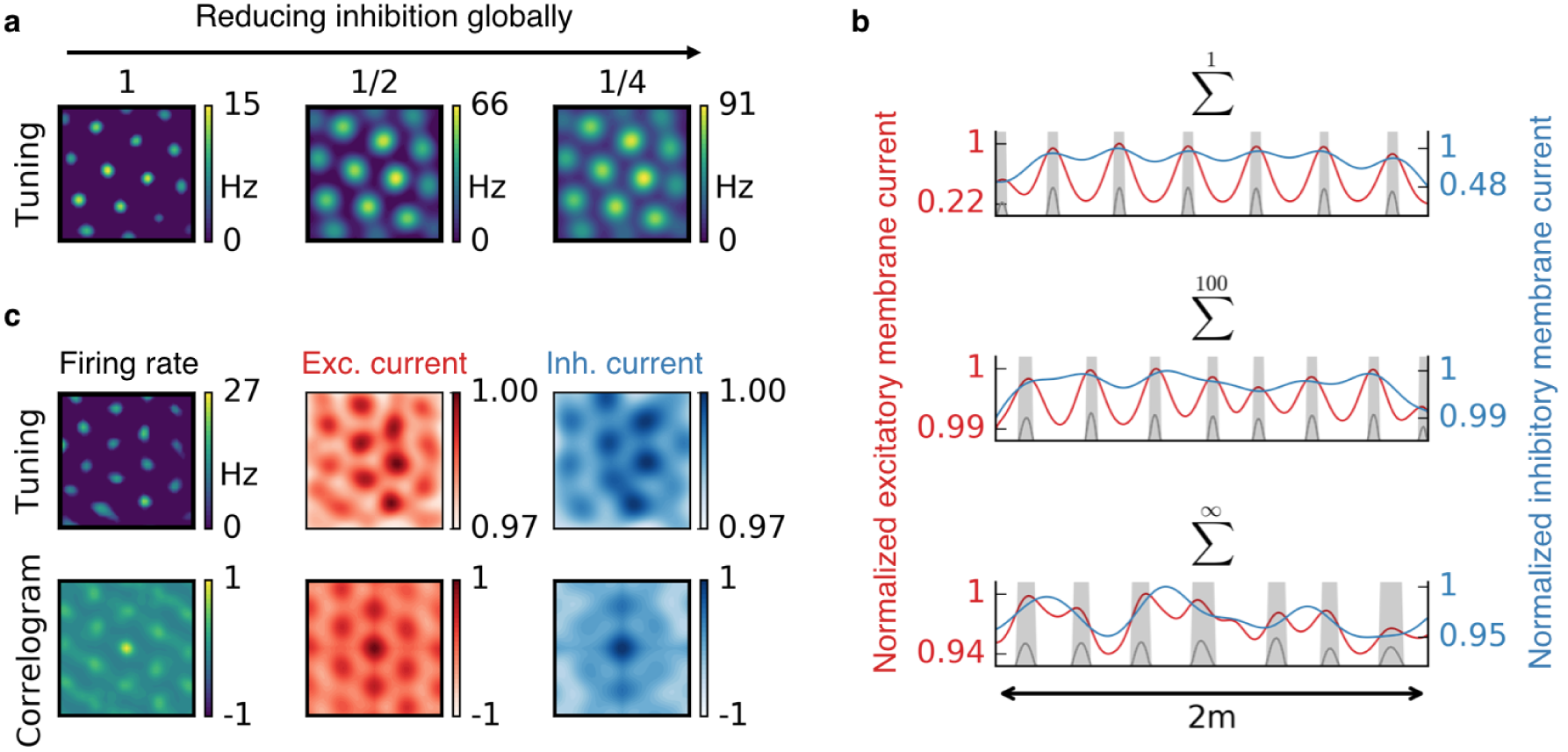
The effect of reduced inhibition on grid cell properties. **a**) Reducing the strength of inhibitory synapses to a fraction of its initial value (from left to right: 1, 1/2, 1/4) leads to larger grid fields but an unchanged grid spacing in our model. In continuous attractor network models, the same reduction of inhibition would affect not only the field size but also the grid spacing. **b**) Excitatory (red) and inhibitory (blue) membrane current to a cell with grid like firing pattern (gray) on a linear track. The currents are normalized to a maximum value of 1. Different rows correspond to different spatial tuning characteristics of the input neurons. From top to bottom: Place cell-like tuning, sparse non-localized tuning (sum of 100 randomly located place fields), dense non-localized tuning (Gaussian random fields). Peaks in excitatory membrane current are colocalized with grid fields (shaded area) for all input statistics. In contrast, the inhibitory membrane current is not necessarily correlated with the grid fields for non-localized input. Moreover, the dynamic range of the membrane currents is reduced for non-localized input. A reduction of inhibition as shown in **a** corresponds to a lowering of the blue curve. **c**) Excitatory and inhibitory membrane current to a grid cell receiving sparse non-localized input (sum of 100 randomly located place fields) in two dimensions. Top: Tuning of output firing rate, normalized excitatory and inhibitory membrane current. Bottom: autocorrelograms thereof. The grid pattern is strongly apparent in the spatial tuning of the excitatory membrane current but less in the inhibitory membrane current.

The grid patterns of topologically nearby grid cells in the mEC typically have the same orientation and spacing but different phases [4]. This property is immanent to CAN models. In contrast, single cell models [14 15 17–19] require additional mechanisms, such as recurrent connections [66], to develop a coordination of neighboring grid cells.

Models that learn grid cells from spatially tuned input do not have to assume a preexisting connectivity pattern or specific mechanisms for path integration [14], but are challenged by the fast emergence of hexagonal firing patterns in unfamiliar environments [4]. Most plasticity-based models require slow learning, such that the animal explores the whole arena before significant synaptic changes occur. Therefore, grid patterns typically emerge slower than experimentally observed [18, 62]. This delay is particularly pronounced in models that require an extensive exploration of both space and movement direction [15]. For the mechanism we suggested, the selforganization was very robust and allowed a rapid pattern formation on short time scales, similar to those observed in rodents (Fig. 3). This speed could be further increased by accelerated reactivation of previous experiences during periods of rest [67]. By this means, the exploration time and the time it takes to activate all input patterns could be decoupled, leading to a much faster emergence of grid cells in all trajectory-independent models with associative learning. Other models that explain the emergence of grid patterns from place cell input through synaptic depression and potentiation also develop grid cells in realistic times [17, 19]. How these models generalize to potentially non-localized input is yet to be shown.

Experiments show that the pattern and the orientation of grid cells is influenced by the geometry of the environment. In a quadratic box, the orientation of grid cells tends to align – with a small offset - to one of the box axes [68]. In trapezoidal arenas, the hexagonality of grids is distorted [69]. We considered quadratic and circular (SOM, Fig. **S11**) arenas with rat trajectories from behavioral experiments and found that the boundaries distort the grid pattern also in our simulations, particularly for localized inputs (SOM, Fig. **S11**). In trapezoidal geometries, we expect this to lead to non-hexagonal grids. However, we did not observe a pronounced alignment to quadratic boundaries, if the input place fields are randomly located (SOM, Fig. **S11**).

We found that interacting excitatory and inhibitory plasticity serves as a simple and robust mechanism for rapid self-organization of stable and symmetric patterns from spatially modulated feedforward input. The suggested mechanism ports the robust pattern formation of attractor models [9, 10, 12] from the neural to the spatial domain and increases the speed of self-organization of plasticity-based mechanisms [15, 17-19] to time scales on which the spatial tuning of neurons is typically measured. It will be interesting to explore how recurrent connections between output cells can help to understand the role of local inhibitory connections [13] and the presence or absence of topographic arrangements of spatially tuned cells [59, 70, 71]. We illustrated the properties and requirements of the model in the realm of spatial representations. Since invariance and selectivity are ubiquitous properties of receptive fields in the brain, the interaction of excitatory and inhibitory synaptic plasticity might be essential to form stable representations from sensory input also in other brain areas [39, 72].

## Online Methods

### Network architecture and neuron model

We study a feedforward network where a single output neuron receives synaptic input from *N*_E_ excitatory and *N*_I_ inhibitory neurons (Fig. 1a) with synaptic weight vectors **w**^E^, **w**^I^ and spatially tuned rates **r**^E^(**x**), **r**^I^(**x**), respectively. Here **x** denotes the location and later also the head direction of the animal. For simplicity and to allow a mathematical analysis we use a rate-based description for all neurons. The firing rate of the output neuron is given by the rectified sum of weighted excitatory and inhibitory inputs:

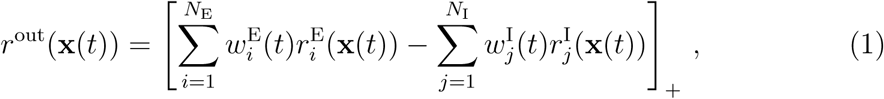

where [∙]_+_ denotes a rectification that sets negative firing rates to zero. To comply with the notion of excitation and inhibition, all weights are constrained to be positive. In most simulations we use four times as many excitatory as inhibitory input neurons (Tables **1** to **3** in the SOM list values for all simulation parameters used in each figure).

### Excitatory and inhibitory plasticity

In each unit time step (Δ*t =* 1), the excitatory weights are updated according to a Hebbian rule:

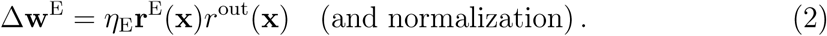

The excitatory learning rate *η*_E_ is a constant that we chose individually for each simulation. To avoid unbounded weight growth, we use a quadratic multiplicative normalization, i.e., we keep the sum of the squared weights of the excitatory population 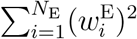 constant at its initial value, by rescaling the weights after each unit time step. We model inhibitory synaptic plasticity using a previously suggested learning rule [37]:

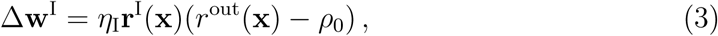

with inhibitory learning rate *η_I_* and target rate *ρ*_0_. We always use *η_I_* > *η_E_* and *ρ_0_* = 1 Hz. Negative inhibitory weights are set to zero.

### Rat trajectory

In the linear track model (one dimension, Figs. 1 and 6), we create artificial run-and-tumble trajectories *x*(*t*) constrained on a line of length *L* with constant velocity *ν* =1 cm per unit time step and persistence length *L*/2 (SOM).

In the open arena model (two dimensions, Figs. 2 to 4 and 6), we use trajectories *x*(*t*) from behavioral data [40] of a rat that moved in a 1m × 1m quadratic enclosure (SOM).

In the model for neurons with head direction tuning (three dimensions, Fig. 5), we use the same behavioral trajectories as in two dimensions and model the head direction as noisily aligned to the direction of motion (SOM).

### Spatially tuned inputs

The firing rates of excitatory and inhibitory synaptic inputs 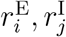 are tuned to the location **x** of the animal. In the following, we use *x* and *y* for the first and second spatial dimension and *z* for the head direction.

For place field-like input, we use Gaussian tuning functions with standard deviation *σ*_E_*, σ*_I_ for the excitatory and inhibitory population, respectively. In Fig. 4 the standard deviation is chosen independently along the *x* and *y* direction. The centers of the Gaussians are drawn randomly from a distorted lattice (SOM). This way we ensure random but spatially dense tuning. The lattice contains locations outside the box to reduce boundary effects.

For sparse non-localized input with 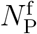 fields per neuron of population P, we first create 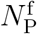 distorted lattices, each with *N*_P_ locations. We then assign 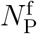 of the resulting 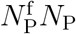 locations at random and without replacement to each input neuron (SOM).

For dense non-localized input, we convolve Gaussians with white noise and increase the resulting signal to noise ratio by setting the minimum to zero and the mean to 0.5 (SOM). The Gaussian convolution kernels have different standard deviations for different populations. For each input neuron we use a different realization of white noise. This results in arbitrary tuning functions of the same autocorrelation length as the – potentially asymmetric – Gaussian convolution kernel.

For combined spatially and head direction tuned input, we use the Gaussian tuning curves described above for the spatial tuning and von Mises distributions along the head direction dimension.

### Initial synaptic weights and global reduction of inhibition

We specify a mean for the initial excitatory and inhibitory weights, respectively, and randomly draw each synaptic weight from the corresponding mean ±5%. The excitatory mean is chosen such that the output neuron would fire above the target rate everywhere in the absence of inhibition; we typically take this mean to be 1 (Table **1**, SOM). The mean inhibitory weight is then determined such that the output neuron would fire close to the target rate, if all the weights were at their mean value (Table **2**, SOM).

We modeled a global reduction of inhibition by scaling all inhibitory weights by a constant factor, after the grid has been learned.

## Acknowledgments

We are grateful to W. Gerstner for feedback on the manuscript and O. Mackwood for providing software tools. This work was funded by the German Federal Ministry for Education and Research, FKZ 01GQ1201.

## References

[1] J. O’Keefe, “Place units in the hippocampus of the freely moving rat,” Experimental neurology, vol. 51, no. 1, pp. 78–109, (1976).

[2] J. S. Taube, R. U. Muller, and J. B. Ranck, “Head-direction cells recorded from the postsubiculum in freely moving rats. ii. effects of environmental manipulations,” The Journal of Neuroscience, vol. 10, no. 2, pp. 436–447, (1990).

[3] M. Fyhn, S. Molden, M. P. Witter, E. I. Moser, and M.-B. Moser, “Spatial representation in the entorhinal cortex,” Science, vol. 305, no. 5688, pp. 1258–1264, (2004).

[4] T. Hafting, M. Fyhn, S. Molden, M.-B. Moser, and E. I. Moser, “Microstructure of a spatial map in the entorhinal cortex,” Nature, vol. 436, no. 7052, pp. 801806, (2005).

[5] R. U. Muller, E. Bostock, J. S. Taube, and J. L. Kubie, “On the directional firing properties of hippocampal place cells,” The Journal of Neuroscience, vol. 14, no. 12, pp. 7235–7251, (1994).

[6] N. Burgess, F. Cacucci, C. Lever, and J. O’Keefe, “Characterizing multiple independent behavioral correlates of cell firing in freely moving animals,” Hippocampus, vol. 15, no. 2, pp. 149–153, (2005).

[7] F. Sargolini, M. Fyhn, T. Hafting, B. L. McNaughton, M. P. Witter, M.-B. Moser, and E. I. Moser, “Conjunctive representation of position, direction, and velocity in entorhinal cortex,” Science, vol. 312, no. 5774, pp. 758–762, (2006).

[8] J. Krupic, N. Burgess, and J. O’Keefe, “Neural representations of location composed of spatially periodic bands,” Science, vol. 337, no. 6096, pp. 853–857, (2012).

[9] M. C. Fuhs and D. S. Touretzky, “A spin glass model of path integration in rat medial entorhinal cortex,” The Journal of Neuroscience, vol. 26, no. 16, pp. 4266–4276, (2006).

[10] B. L. McNaughton, F. P. Battaglia, O. Jensen, E. I. Moser, and M.-B. Moser, “Path integration and the neural basis of the’cognitive map’,” Nature Reviews Neuroscience, vol. 7, no. 8, pp. 663–678, (2006).

[11] M. Franzius, H. Sprekeler, and L. Wiskott, “Slowness and sparseness lead to place, head-direction, and spatial-view cells,” PLoS Comput Biol, vol. 3, no. 8, p. e166, (2007).

[12] Y. Burak and I.R. Fiete, “Accurate path integration in continuous attractor network models of grid cells,” PLoS Comput Biol, vol. 5, no. 2, p. e1000291, (2009).

[13] J.J. Couey, A. Witoelar, S.-J. Zhang, K. Zheng, J. Ye, B. Dunn, R. Czajkowski, M.-B. Moser, E.I. Moser, Y. Roudi, et al., “Recurrent inhibitory circuitry as a mechanism for grid formation,” Nature neuroscience, vol. 16, no. 3, pp. 318–324, (2013).

[14] N. Burgess, C. Barry, and J. O’Keefe, “An oscillatory interference model of grid cell firing,” Hippocampus, vol. 17, no. 9, pp. 801–812, (2007).

[15] E. Kropff and A. Treves, “The emergence of grid cells: Intelligent design or just adaptation?,” Hippocampus, vol. 18, no. 12, pp. 1256–1269, (2008).

[16] D. Bush and N. Burgess, “A hybrid oscillatory interference/continuous attractor network model of grid cell firing,” The Journal of Neuroscience, vol. 34, no. 14, pp. 5065–5079, (2014).

[17] L. Castro and P. Aguiar, “A feedforward model for the formation of a grid field where spatial information is provided solely from place cells,” Biological cybernetics, vol. 108, no. 2, pp. 133–143, (2014).

[18] Y. Dordek, D. Soudry, R. Meir, and D. Derdikman, “Extracting grid cell characteristics from place cell inputs using non-negative principal component analysis,” eLife, vol. 5, p. e10094, (2016).

[19] A. Stepanyuk, “Self-organization of grid fields under supervision of place cells in a neuron model with associative plasticity,” Biologically Inspired Cognitive Architectures, vol. 13, pp. 48–62, (2015).

[20] L.M. Giocomo, M.-B. Moser, and E.I. Moser, “Computational models of grid cells,” Neuron, vol. 71, no. 4, pp. 589–603, (2011).

[21] E.A. Zilli, “Models of grid cell spatial firing published 2005–2011,” Frontiers in neural circuits, vol. 6, p. 16, (2012).

[22] M. Tsodyks and T. Sejnowski, “Associative memory and hippocampal place cells,” International journal of neural systems, vol. 6, pp. 81–86, (1995).

[23] F.P. Battaglia and A. Treves, “Attractor neural networks storing multiple space representations: a model for hippocampal place fields,” Physical Review E, vol. 58, no. 6, p. 7738, (1998).

[24] A. Arleo and W. Gerstner, “Spatial cognition and neuro-mimetic navigation: a model of hippocampal place cell activity,” Biological cybernetics, vol. 83, no. 3, pp. 287–299, (2000).

[25] T. Solstad, E.I. Moser, and G.T. Einevoll, “From grid cells to place cells: a mathematical model,” Hippocampus, vol. 16, no. 12, pp. 1026–1031, (2006).

[26] M. Franzius, R. Vollgraf, and L. Wiskott, “From grids to places,” Journal of computational neuroscience, vol. 22, no. 3, pp. 297–299, (2007).

[27] N. Burgess and J. O’Keefe, “Models of place and grid cell firing and theta rhythmicity,” Current opinion in neurobiology, vol. 21, no. 5, pp. 734–744, (2011).

[28] B. McNaughton, L. Chen, and E. Markus, ““dead reckoning,” landmark learning, and the sense of direction: a neurophysiological and computational hypothesis,” Journal of Cognitive Neuroscience, vol. 3, no. 2, pp. 190–202, (1991).

[29] A.D. Redish, A.N. Elga, and D.S. Touretzky, “A coupled attractor model of the rodent head direction system,” Network: Computation in Neural Systems, vol. 7, no. 4, pp. 671–685, (1996).

[30] K. Zhang, “Representation of spatial orientation by the intrinsic dynamics of the head-direction cell ensemble: a theory,” The journal of neuroscience, vol. 16, no. 6, pp. 2112–2126, (1996).

[31] J.J. DiCarlo and D.D. Cox, “Untangling invariant object recognition,” Trends in cognitive sciences, vol. 11, no. 8, pp. 333–341, (2007).

[32] C. Buetfering, K. Allen, and H. Monyer, “Parvalbumin interneurons provide grid cell-driven recurrent inhibition in the medial entorhinal cortex,” Nature neuroscience, vol. 17, no. 5, pp. 710–718, (2014).

[33] J. O’Keefe and J. Dostrovsky, “The hippocampus as a spatial map. preliminary evidence from unit activity in the freely-moving rat,” Brain research, vol. 34, no. 1, pp. 171–175, (1971).

[34] L. Marshall, D.A. Henze, H. Hirase, X. Leinekugel, G. Dragoi, and G. Buzsaki, “Hippocampal pyramidal cell-interneuron spike transmission is frequency dependent and responsible for place modulation of interneuron discharge,” J Neurosci, vol. 22, no. 2, (2002).

[35] W.B. Wilent and D.A. Nitz, “Discrete place fields of hippocampal formation interneurons,” Journal of neurophysiology, vol. 97, no. 6, pp. 4152–4161, (2007).

[36] D.O. Hebb, The organization of behavior: A neuropsychological approach. John Wiley & Sons, (1949).

[37] T. Vogels, H. Sprekeler, F. Zenke, C. Clopath, and W. Gerstner, “Inhibitory plasticity balances excitation and inhibition in sensory pathways and memory networks,” Science, vol. 334, no. 6062, pp. 1569–1573, (2011).

[38] T. Hafting, M. Fyhn, T. Bonnevie, M.-B. Moser, and E.I. Moser, “Hippocampus-independent phase precession in entorhinal grid cells,” Nature, vol. 453, no. 7199, pp. 1248–1252, (2008).

[39] C. Clopath, T.P. Vogels, R.C. Froemke, and H. Sprekeler, “Receptive field formation by interacting excitatory and inhibitory synaptic plasticity,” bioRxiv, p. 066589, (2016).

[40] F. Sargolini, M. Fyhn, T. Hafting, B.L. McNaughton, M.P. Witter, M.-B. Moser, and E.I. Moser, “http://www.ntnu.edu/kavli/research/grid-cell-data,” (2016).

[41] A.M. Turing, “The chemical basis of morphogenesis,” Philosophical Transactions of the Royal Society of London B: Biological Sciences, vol. 237, no. 641, pp. ST-72, (1952).

[42] M. Fyhn, T. Hafting, A. Treves, M.-B. Moser, and E.I. Moser, “Hippocampal remapping and grid realignment in entorhinal cortex,” Nature, vol. 446, no. 7132, pp. 190–194, (2007).

[43] F. Savelli, D. Yoganarasimha, and J.J. Knierim, “Influence of boundary removal on the spatial representations of the medial entorhinal cortex,” Hippocampus, vol. 18, no. 12, pp. 1270–1282, (2008).

[44] C. Barry, L.L. Ginzberg, J. O’Keefe, and N. Burgess, “Grid cell firing patterns signal environmental novelty by expansion,” Proceedings of the National Academy of Sciences, vol. 109, no. 43, pp. 17687–17692, (2012).

[45] E.I. Moser, E. Kropff, and M.-B. Moser, “Place cells, grid cells, and the brain’s spatial representation system,” Neuroscience, vol. 31, no. 1, p. 69, (2008).

[46] S. Leutgeb, J.K. Leutgeb, C.A. Barnes, E.I. Moser, B.L. McNaughton, and M.-B. Moser, “Independent codes for spatial and episodic memory in hippocampal neuronal ensembles,” Science, vol. 309, no. 5734, pp. 619–623, (2005).

[47] J.S. Taube, “Head direction cells recorded in the anterior thalamic nuclei of freely moving rats,” The Journal of neuroscience, vol. 15, no. 1, pp. 70–86, (1995).

[48] L.L. Chen, L.H. Lin, C.A. Barnes, and B.L. McNaughton, “Head-direction cells in the rat posterior cortex. ii. contributions of visual and ideothetic information to the directional firing.,” Experimental brain research, vol. 101, no. 1, pp. 24–34, (1993).

[49] A. Rubin, M.M. Yartsev, and N. Ulanovsky, “Encoding of head direction by hippocampal place cells in bats,” The Journal of Neuroscience, vol. 34, no. 3, pp. 1067–1080, (2014).

[50] J.A. D’amour and R.C. Froemke, “Inhibitory and excitatory spike-timing-dependent plasticity in the auditory cortex,” Neuron, vol. 86, no. 2, pp. 514–528, (2015).

[51] A. Wertz, S. Trenholm, K. Yonehara, D. Hillier, Z. Raics, M. Leinweber, G. Szalay, A. Ghanem, G. Keller, B. Rozsa, et al., “Single-cell-initiated monosynaptic tracing reveals layer-specific cortical network modules,” Science, vol. 349, no. 6243, pp. 70–74, (2015).

[52] D.E. Wilson, D.E. Whitney, B. Scholl, and D. Fitzpatrick, “Orientation selectivity and the functional clustering of synaptic inputs in primary visual cortex,” Nature neuroscience, (2016).

[53] N.M. Van Strien, N. Cappaert, and M.P. Witter, “The anatomy of memory: an interactive overview of the parahippocampal-hippocampal network,” Nature Reviews Neuroscience, vol. 10, no. 4, pp. 272–282, (2009).

[54] R.F. Langston, J.A. Ainge, J.J. Couey, C.B. Canto, T.L. Bjerknes, M.P. Witter, E.I. Moser, and M.-B. Moser, “Development of the spatial representation system in the rat,” Science, vol. 328, no. 5985, pp. 1576–1580, (2010).

[55] T.J. Wills, F. Cacucci, N. Burgess, and J. O’Keefe, “Development of the hippocampal cognitive map in preweanling rats,” Science, vol. 328, no. 5985, pp. 1573–1576, (2010).

[56] T. Bonnevie, B. Dunn, M. Fyhn, T. Hafting, D. Derdikman, J.L. Kubie, Y. Roudi, E.I. Moser, and M.-B. Moser, “Grid cells require excitatory drive from the hippocampus,” Nature neuroscience, vol. 16, no. 3, pp. 309–317, (2013).

[57] M.W. Jung, S.I. Wiener, and B.L. McNaughton, “Comparison of spatial firing characteristics of units in dorsal and ventral hippocampus of the rat,” The Journal of neuroscience, vol. 14, no. 12, pp. 7347–7356, (1994).

[58] V.H. Brun, T. Solstad, K.B. Kjelstrup. M. Fyhn, M.P. Witter, E.I. Moser, and M.-B. Moser, “Progressive increase in grid scale from dorsal to ventral medial entorhinal cortex,” Hippocampus, vol. 18, no. 12, pp. 1200–1212, (2008).

[59] H. Stensola, T. Stensola, T. Solstad, K. Frøland, M.-B. Moser, and E.I. Moser, “The entorhinal grid map is discretized,” Nature, vol. 492, no. 7427, pp. 72–78, (2012).

[60] S. Melzer, M. Michael, A. Caputi, M. Eliava, E.C. Fuchs, M.A. Whittington, and H. Monyer, “Long-range-projecting gabaergic neurons modulate inhibition in hippocampus and entorhinal cortex,” Science, vol. 335, no. 6075, pp. 1506–1510, (2012).

[61] P. Beed, A. Gundlfinger, S. Schneiderbauer, J. Song, C. Böhm, A. Burgalossi, M. Brecht, I. Vida, and D. Schmitz, “Inhibitory gradient along the dorsoventral axis in the medial entorhinal cortex,” Neuron, vol. 79, no. 6, pp. 1197–1207, (2013).

[62] J. Widloski and I.R. Fiete, “A model of grid cell development through spatial exploration and spike time-dependent plasticity,” Neuron, vol. 83, no. 2, pp. 481495, (2014).

[63] J. Widloski and I.R. Fiete, “Cortical microcircuit determination through global perturbation and sparse sampling in grid cells,” bioRxiv, p. 019224, (2015).

[64] N.I. Cilz, L. Kurada, B. Hu, and S. Lei, “Dopaminergic modulation of gabaergic transmission in the entorhinal cortex: concerted roles of *a1* adrenoreceptors, inward rectifier k+, and t-typ. ca2+ channels,” Cerebral Cortex, p. bht177, (2013).

[65] C. Schmidt-Hieber and M. Häusser, “Cellular mechanisms of spatial navigation in the medial entorhinal cortex,” Nature neuroscience, vol. 16, no. 3, pp. 325331, (2013).

[66] B. Si, E. Kropff, and A. Treves, “Grid alignment in entorhinal cortex,” Biological cybernetics, vol. 106, no. 8–9, pp. 483–506, (2012).

[67] A.K. Lee and M.A. Wilson, “Memory of sequential experience in the hippocampus during slow wave sleep,” Neuron, vol. 36, no. 6, pp. 1183–1194, (2002).

[68] T. Stensola, H. Stensola, M.-B. Moser, and E.I. Moser, “Shearing-induced asymmetry in entorhinal grid cells,” Nature, vol. 518, no. 7538, pp. 207–212, (2015).

[69] J. Krupic, M. Bauza, S. Burton, C. Barry, and J. O’Keefe, “Grid cell symmetry is shaped by environmental geometry,” Nature, vol. 518, no. 7538, pp. 232–235, (2015).

[70] J. O’Keefe, N. Burgess, J.G. Donnett, K.J. Jeffery, and E.A. Maguire, “Place cells, navigational accuracy, and the human hippocampus,” Philosophical Transactions of the Royal Society of London B: Biological Sciences, vol. 353, no. 1373, pp. 1333–1340, (1998).

[71] L.M. Giocomo, T. Stensola, T. Bonnevie, T. Van Cauter,M.-B. Moser, and E.I. Moser, “Topography of head direction cells in medial entorhinal cortex,” Current Biology, vol. 24, no. 3, pp. 252–262, (2014).

[72] A.O. Constantinescu, J.X. O’Reilly, and T.E. Behrens, “Organizing conceptual knowledge in humans with a gridlike code,” Science, vol. 352, no. 6292, pp. 1464–1468, (2016).

